# TRACT revisited: an algebraic solution for determining overall rotational correlation times from cross-correlated relaxation rates

**DOI:** 10.1101/2021.01.16.426977

**Authors:** Scott A. Robson, Çağdaş Dağ, Hongwei Wu, Joshua J. Ziarek

## Abstract

Accurate rotational correlation times (*τ*_*c*_) are critical for quantitative analysis of fast timescale NMR dynamics. As molecular weights increase, the classic derivation of *τ*_*c*_ using transverse and longitudinal relaxation rates becomes increasingly unsuitable due to the non-trivial contribution of remote dipole-dipole interactions to longitudinal relaxation. Derivations using cross-correlated relaxation experiments, such as TRACT, overcome these limitations but are erroneously calculated in 65% of the citing literature. Herein, we developed an algebraic solutions to the Goldman relationship that facilitate rapid, point-by-point calculations for straightforward identification of appropriate spectral regions where global tumbling is likely to be dominant. The rigid-body approximation of the Goldman relationship has been previously shown to underestimate TRACT-based rotational correlation time estimates. This motivated us to develop a second algebraic solution that employs a simplified model-free spectral density function including an order parameter term that could, in principle, be set to an average backbone S^2^ ≈ 0.9 to further improve the accuracy of *τ*_*c*_ estimation. These solutions enabled us to explore the boundaries of the Goldman relationship as a function of the H-N internuclear distance (*r*), difference of the two principal components of the axially-symmetric ^15^N CSA tensor (*Δδ*_*N*_), and angle of the CSA tensor relative to the N-H bond vector (*θ*). We hope our algebraic solutions and analytical strategies will increase the accuracy and application of the TRACT experiment.

## Introduction

A particle’s rotational Brownian diffusion is characterized by the average time to rotate one radian, also known as the rotational correlation time (*τ*_*c*_). It is related to the size and shape of a molecule, and in the case of a rigid, spherical particle, can be estimated from the Stokes-Einstein relation^1^. The rotational correlation time is frequently used in biophysics to gauge molecular aggregation and solvent viscosity. In protein NMR, rotational correlation time estimates are used to optimize interscan recycling delays, magnetization transfer delays in correlation experiments, and indirect dimension evolution times in multidimensional experiments^2^. Perhaps most significantly, *τ*_*c*_ is the critical parameter for quantitative dynamics analyses, such as ‘model-free’ formalism, in which separation of overall and internal motion are required^3,4^.

The 1989 seminal work by Kay, Torchia and Bax^5^ showed *τ*_*c*_ can be estimated from ^15^N longitudinal (R_1_) and transverse (R_2_) relaxation rates using equations established by Abragam^6^. These equations assume there is no rapid internal motion of the internuclear bond vector (i.e. motion faster than the rate of molecular tumbling), and that relaxation only results from i) dipole coupling with the covalently bonded nucleus and ii) the chemical shift anisotropy of the relaxing nucleus. In the original model free formalism, the amplitude and time parameter for this fast internal motion are referred to as the order parameter (S^2^) and the effective correlation time (*τ*_*e*_), respectively, where it is assumed that *τ*_*e*_ ≪ *τ*_*c*_^3,4^. Note that *τ*_*m*_ is frequently used in the literature to refer to overall molecular tumbling time, where 1/*τ*_*m*_ = 1/*τ*_*e*_ + 1/*τ*_*c*_. In this present work, unless stated otherwise, we assume fast motions on the time scale of *τ*_*e*_ ≪ *τ*_*c*_ do not exist and thus *τ*_*c*_ is synonymous with *τ*_*m*_. In the absence of *τ*_*e*_, the S^2^ order parameter essentially cancels out when R_2_ is divided by R_1_^5^. Thus, the R_2_/R_1_ ratio can be used to estimate *τ*_*c*_ by using the approximate formula in Eqn. 1, where V_N_ is the ^15^N resonance frequency (in Hz).

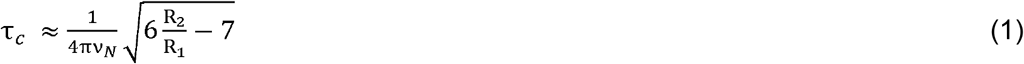

Although this is a common approach for estimating the rotational correlation time, there are several important considerations. First, R_1_ values are susceptible to rapid internal motions of the spin system bond vector; Kay and co-workers established that this could be resolved by excluding all spin systems with ^15^N{^1^H}-NOE values < 0.6^5^. Second, substantial chemical exchange contributions can potentiate the observed R_2_ rates^7^. Again, Kay et al. established a precedence for manually inspecting each ^15^N rate constant for indications of fast internal motions and chemical exchange^5^. This method of excluding spin systems from *τ*_*c*_ estimates was subsequently automated by Clore et al^8^, who first suggested determination of the mean R_2_/R_1_ ratio and exclusion of all values outside one standard deviation. Implicit in this model is that the tumbling is isotropic^5,8^; Barbato et al^9^ expanded application to anisotropic biomolecular systems. Anisotropic tumbling deviates R_2_ and R_1_ values, independent of chemical exchange or fast internal motions, but in opposing directions. Therefore, exchanging residues in which the R_2_/R_1_ model does not apply can be detected by considering R_2_ values that deviate more than one standard deviation from the mean, but without an associated decrease in R_1_ relaxation ^9^. Again, spin systems with ^15^N{^1^H}-NOE < 0.6 (indicating significant fast time scale motion) were excluded from *τ*_*c*_ estimation. The remaining R_2_/R_1_ values are then used to calculate *τ*_*c*_ using Eqn 1. Statistical selection of spin systems for *τ*_*c*_ calculation is generally appropriate but does have the potential for mishandling by an inexperienced user. For example, a protein with significant regions undergoing chemical exchange will report high R_2_ or R_2_/R_1_ standard deviations from inclusion of unsuitable spin systems. It is also not hard to imagine systems with significant regions of high internal motions, but not complete random coil, which would skew the distribution of R_1_ values and lead to the inclusion of spin systems where the R_2_/R_1_ model is a poor approximation. Lastly, the R_2_/R_1_ method does not account for the effects of dipolar couplings to remote (non-covalently bonded) protons on R_1_, which increase near exponentially as a function of molecular weight^10,11^; although perdeuteration largely ameliorates this problem. Nonetheless, there are clear advantages to approaches that can estimate *τ*_*c*_ from a measurable relaxation parameter that is insensitive to the effects of remote dipolar couplings and chemical exchange.

One such quantifiable phenomenon is transverse cross-correlated relaxation (CCR, η_*xy*_). CCR results from the coordinated rotation of two nuclei in a magnetic field^12^ and is primarily a function of dipole-dipole (DD) coupling, the chemical shift anisotropy (CSA) of the observed relaxing nucleus, and the molecular rotational correlation time. Measurements of CCR exploit the fact that the sign (±) of *η*_*xy*_ depends on the spin states of the coupled nuclei. Cross-correlated relaxation only contributes to R_2_ and R_1_ in coupled systems because the opposing (opposite sign) contributions to relaxation cancel for decoupled spins^12^. ^1^H-^15^N and ^1^H-^13^C aromatic TROSY experiments exploit this property by only selecting signals from the spin state with relaxation interference^13,14^. Several methods have been developed to measure *η*_*xy*_ in the ^1^H-^15^N spin system^15–18^ where the most common approach is a pair of spectra that record the transverse relaxation rates of ^15^N alpha (R_α_) and beta (R_β_) spin states. These rates are the sum of the auto-relaxation rate (R_auto_), remote ^1^H dipolar interactions (R_D_), chemical exchange (R_ex_), and *η*_*xy*_ (Eqns. 2 and 3); where *η*_*xy*_ is derived from subtraction of R_α_ and R_β_^11,16,17^ (Eqns. 4 and 5).

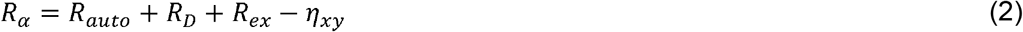

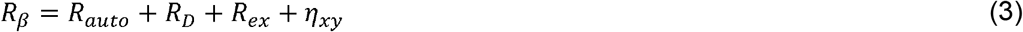

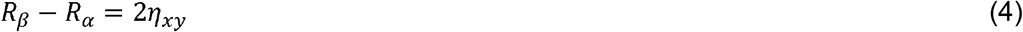

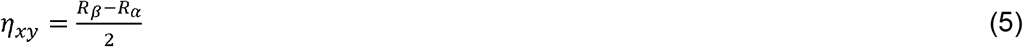

Using this method, the contributions to relaxation from remote protons and chemical exchange are cancelled and extraction of *η*_*xy*_ is possible. Goldman^12^ showed that *η*_*xy*_ can be estimated given *τ*_*c*_, via the spectral density function, using Eqn. 6.

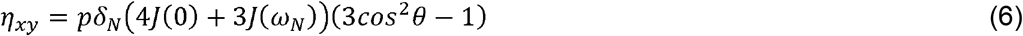

where:

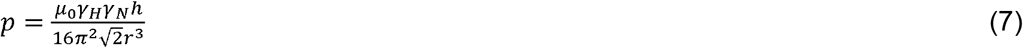

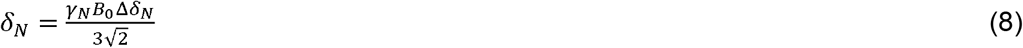

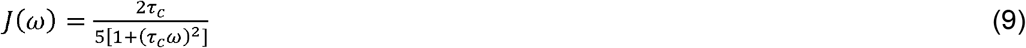

and:

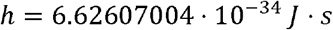

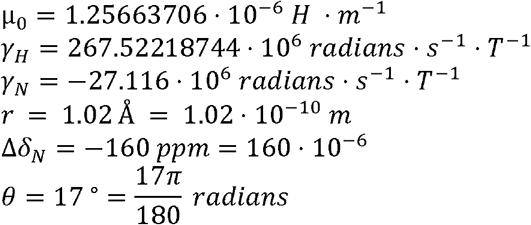

Goldman’s relationship between *η*_*xy*_ and *τ*_*c*_ was first exploited experimentally by Lee and colleagues in the [^15^N,^1^H]-TRACT (<u>T</u>ROSY for <u>r</u>ot<u>a</u>tional <u>c</u>orrelation <u>t</u>imes) pulse sequence^11^; although, the manuscript does not explicitly detail how the Goldman relationship was solved. There are three important assumptions when applying the Goldman approach to TRACT data: i) the region analyzed is free of fast internal motions, which is rarely known *a priori*, ii) the system is a rigid body (i.e. S^2^ ~ 1.0), and iii) the spin pair is isolated from remote DD/CSA relaxation interference. When pursuing our own TRACT analyses, we noted our *τ*_*c*_ calculations were inconsistent with the original manuscript despite using identical physical and geometric constants.

Herein, we report an algebraic solution to the Goldman approach for straightforward calculation of *τ*_*c*_ from measured R_α_ and R_β_ relaxation rate. Using this solution, we show that a numerical error in the original TRACT report has propagated into a high proportion of the citing literature. We use our algebraic solution to investigate complications from fast internal motions and propose analytical strategies to exclude unsuitable spin systems. The impact of order parameters motivated us to develop a second algebraic solution that includes S^2^ as a parameter. We also noted that little attention has been paid to distributions of the difference of the two principal components of the axially-symmetric CSA tensor (*Δδ*_*N*_), the CSA tensor angle relative to the internuclear bond vector (θ), and the internuclear distance (*r*). We show that the chosen value can have a non-negligible effect on *τ*_*c*_ calculations but, as the relationship is near linear, a symmetrical random distribution around an average value would cancel out over many spin systems. Finally, we discuss how relaxation interference between remote ^1^H DD and local CSA affects the observed CCR rate; a phenomenon that is independent of R_D_ above but is generally negligible compared to other factors discussed in this paper.

### Experimental

Uniformly-labeled [*U*-^15^N,^2^H]-OmpX was expressed, purified, and solubilized into 0.5% (w/v) dodecylphosphocholine (DPC) micelles as previously described^19^. Final buffer conditions were: 20 mM NaPi (pH 6.8), 100 mM NaCl, 5 mM EDTA, and 10% D_2_O. NMR experiments were performed at 303.15 K on Varian 800 MHz spectrometer equipped with Agilent 5 mm PFG ^1^H{^13^C,^15^N} triple resonance salt tolerant cold probes. 1D TRACT experiments were collected with 4096 complex points and 1.5 s relaxation delay. A series of experiments were collected with eight variable relaxation delays: 1, 2, 4, 8, 16, 32, 64, and 128 ms. Relaxation rates were determined by fitting to a two-parameter exponential function.

## Results and Discussion

### An algebraic solution to the Goldman relationship

While validating our own numerical solution to the Goldman relationship, we noted a 6.6% overestimation of η_xy_ in Lee et al^11^. Specifically, Figures 3 and 4 in their paper present *τ*_*c*_ = 21 ns and 24 ns, respectively, from which we calculate *η*_*xy*_ = 27.1 Hz and 30.9 Hz, respectively, using Eqns. 6-9 and the quoted physical constants. This is inconsistent with the reported *R*_*α*_ and *R*_*β*_, which yield *η*_*xy*_ rates of (64-13)/2 = 25.5 Hz and (80-22)/2 = 29 Hz, respectively, using Eqns. 4-5. Hypothesizing that this discrepancy may be the result of poor numerical minimization, we generated an exact solution to Eqn. 6 with respect to *τ*_*c*_, given B_0_ and *η*_*xy*_.

**Figure 1:**
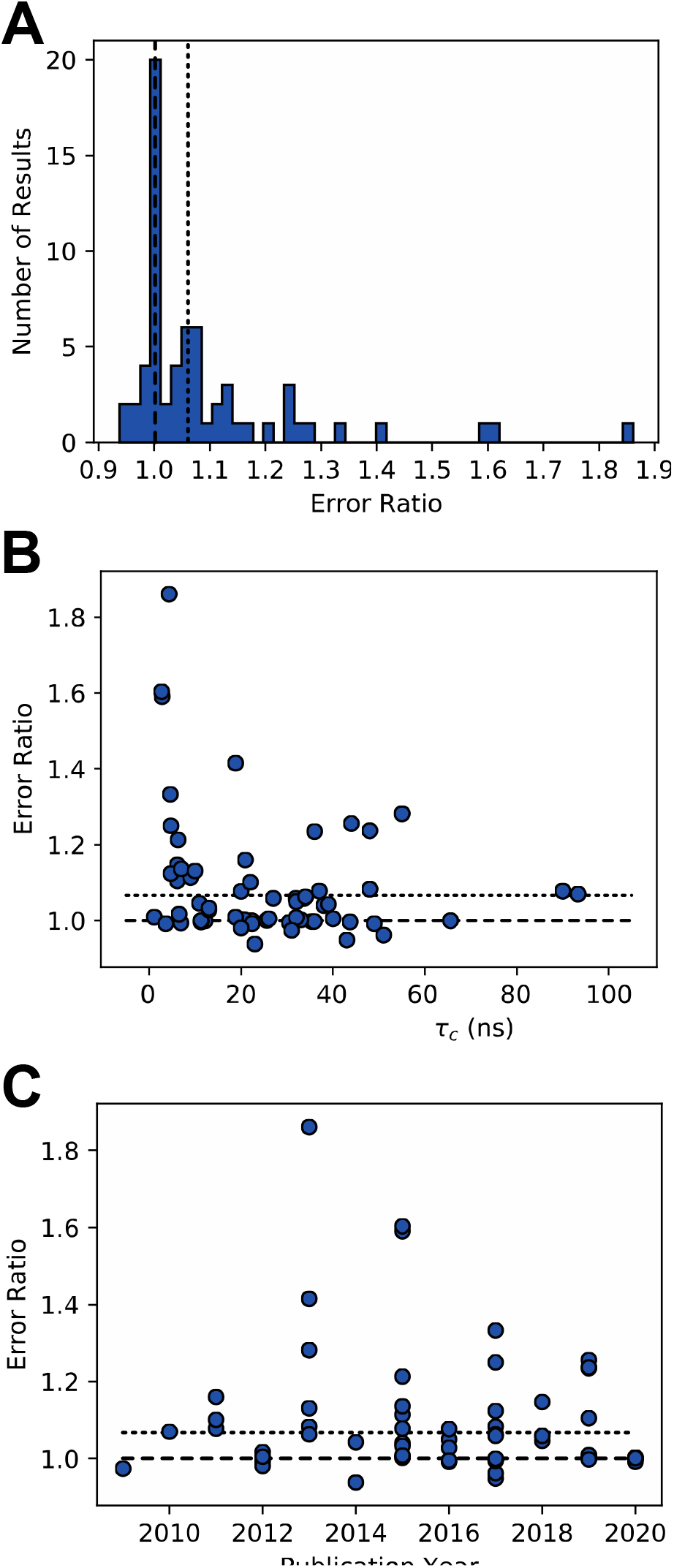
Error analysis of 65 published TRACT-derived rotational correlation times (*τ*_*c*_). The error ratio is defined as the published *τ*_*c*_ divided by the algebraically-determined value using Eqn 12. In all panels, the dashed line (−) and dotted line (…) denote error ratios of 1.0 and 1.067, respectively. A) Histogram of error ratios reveals two clusters: 35% (23/65) of results are narrowly distributed around the accurate result at 1.0 (dashed line), and 23% (15/65) of results are centered at an error ratio of ~1.067 (dotted line) with a slightly wider distribution. The remaining 29% (19/65) of results tend to be overestimates and do not cluster. B) Scatter plot of errors versus year of publication indicate no clear trend that calculation errors are diminishing. C) A scatter plot of *τ*_*c*_ versus error ratio demonstrating there is generally no trend between *τ*_*c*_ and error ratio, apart from a group of large errors for small *τ*_*c*_ values.

**Figure 2:**
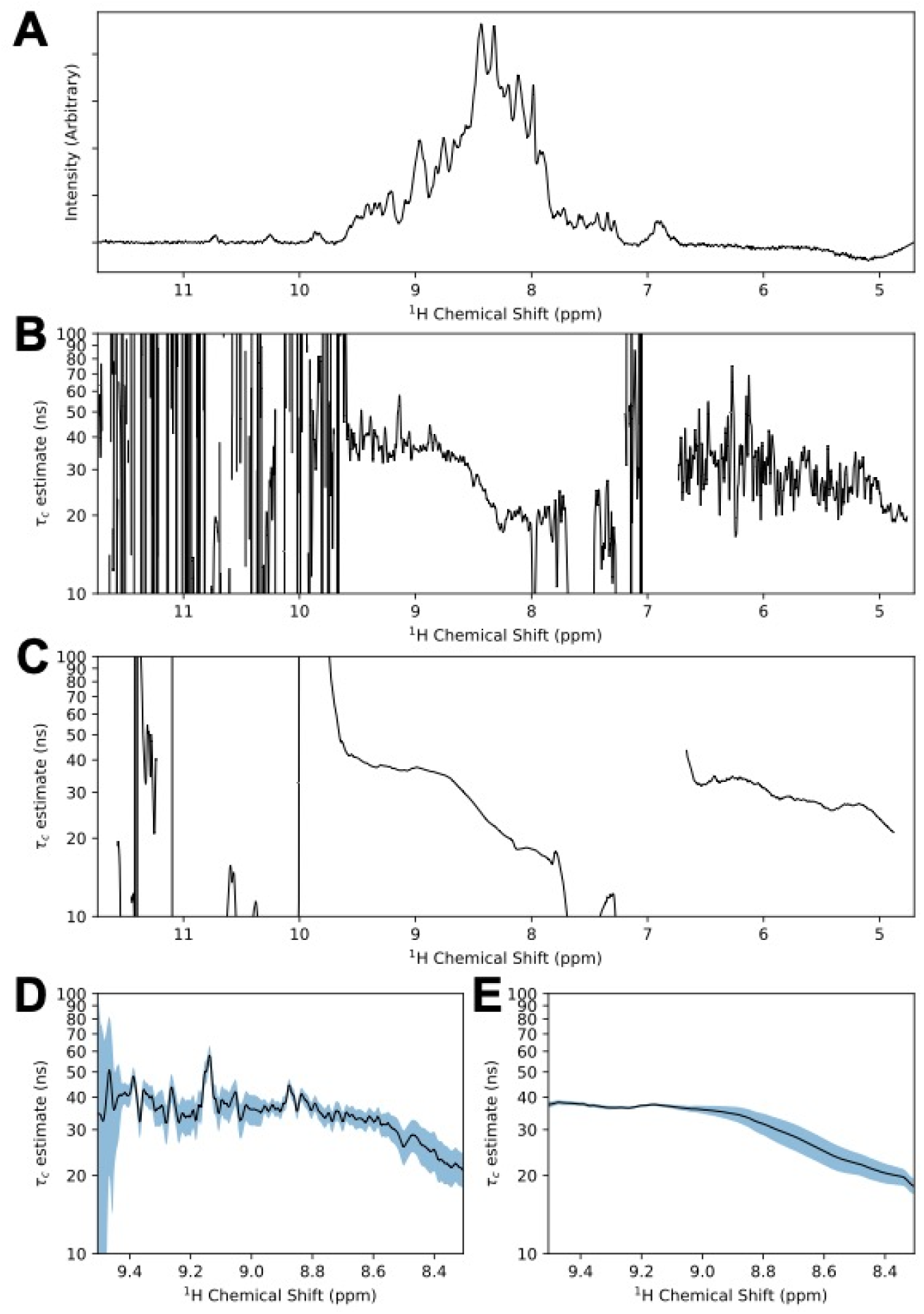
Analysis of TRACT data on OmpX in DPC micelles. A) The amide region (4.70 - 11.74 ppm; 4096 complex points) of the ^15^N-filtered 1D ^1^H_N_ TRACT spectrum for [^15^N,^2^H]-OmpX is reproduced. The spectrum represents the first relaxation delay (1 ms) for the TROSY component (i.e. N_α_ spin state). B) Point-by-point estimations of the rotational correlation time. Calculated values vary wildly even where signals are intense and dispersed (e.g. 9.5 - 8.8 ppm). C) Estimation of *τ*_*c*_ based on a 200 point sliding window (~5% of 4096 complex points). A region of dispersed signals with consistent τ calculations can now be seen from approximately 8.8 – 9.5 ppm. D,E) Expansion of 8.3 - 9.5 ppm region from panels B and C, respectively. The blue regions represent one standard deviation based on a sampling of 200 points around a given point in the spectrum. Application of the sliding window improves *τ*_*c*_ estimates over the 9.17 – 9.5 ppm region from 37.92 ± 4.25 ns to 37.17 ± 0.59 ns.

**Figure 3:**
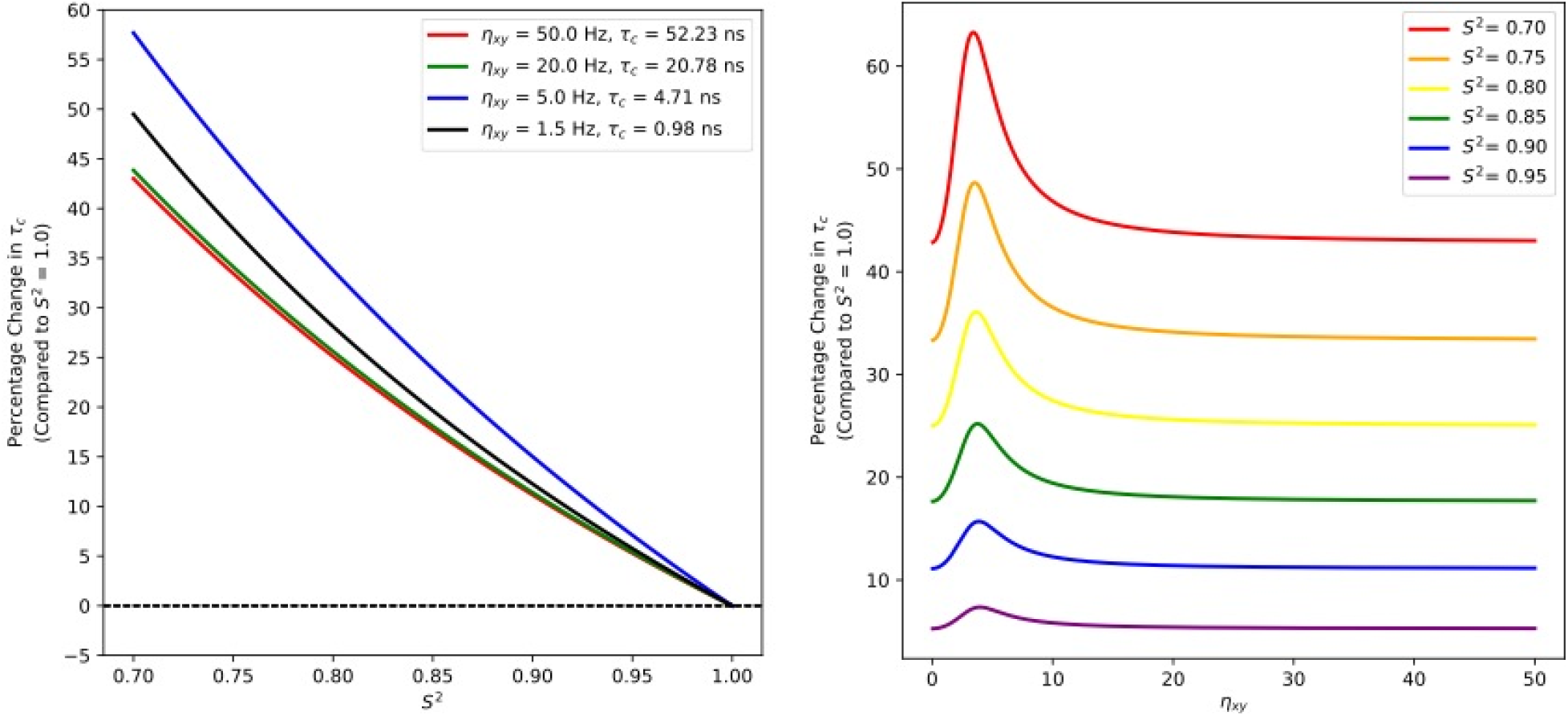
Effect of rigid-body approximation on rotational correlation time estimates derived from modified Goldman relationship. A) Plot of percentage change in *τ*_*c*_ estimation, relative to rigid body approximation, as a function of S^2^ order parameter. Relative to the rigid body assumption, typical backbone order parameters (0.85 ≤ S^2^ ≤ 0.95)^23,24^ increase the rotational correlation time 5-25% depending on the CCR rate; this error increases to ≥45% at an S^2^ ≈ 0.7. B) Plot of percentage change in τ estimation, relative to rigid body approximation, versus η_*xy*_ for select S^2^ values from 0.7 to 0.95. A sharp, biphasic rise in error is observed for η_*xy*_ ≈ 4 Hz.

**Figure 4:**
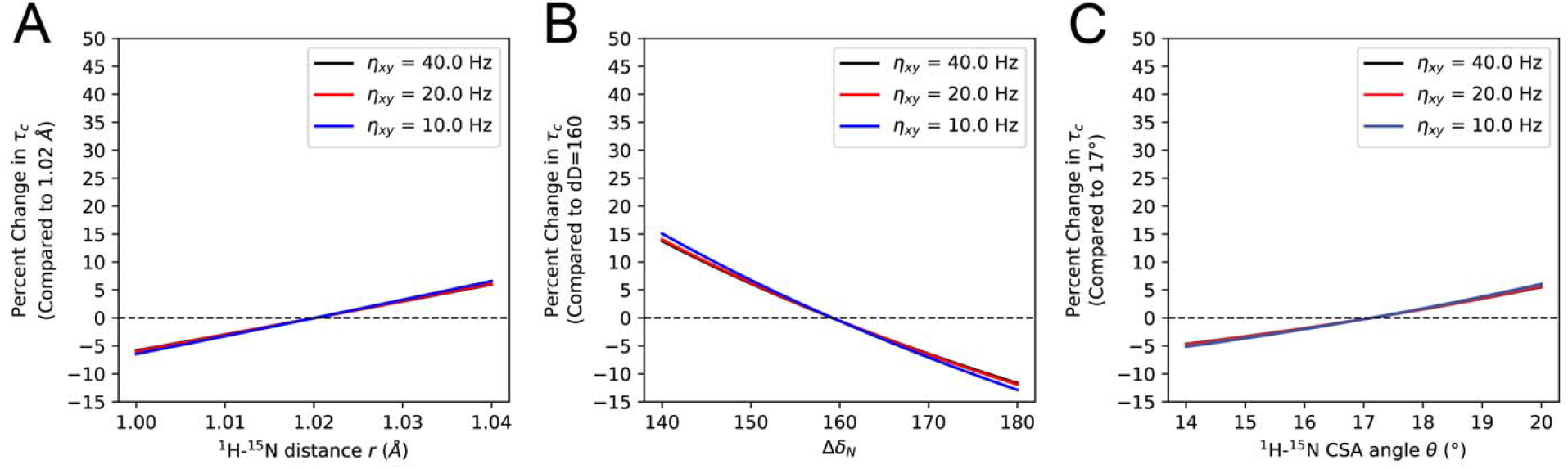
Effect of the internuclear distance (*r*), difference of the two principal components of the axially-symmetric CSA tensor (*Δδ*_*N*_), and angle of the CSA tensor relative to the N-H bond vector (θ) on rotational correlation time *τ*_*c*_ estimation. The relative values simulated are taken from the original TRACT manuscript^11^; as discussed in the text, other values for these parameters may be more accurate. A) Percentage change in *τ*_*c*_ estimation versus *r*, (relative to *r*, = 1.02 Å) for η_*xy*_ ranging from 10 - 40 Hz. B) Percentage change in *τ*_*c*_ estimation versus Δδ_N_ (relative to Δδ_N_ = −160 ppm) for η_*xy*_ ranging from 10 - 40 Hz. C) Percentage change in *τ*_*c*_ estimation versus θ (relative to θ_*xy*_ = 17°) for η_*xy*_ ranging from 10 - 40 Hz. The error for each parameter is negatively symmetric around the chosen value with little variation over the range of simulated η_*xy*_ from 10 to 40 Hz. When integrated over many spins, any deviations around the mean would tend to average out the error in *τ*_*c*_ estimation.

We start by expanding the spectral density function (*J*) in Eqn. 6 and substituting *η*_*xy*_ with (*R*_*β*_ - *R*_*α*_)/2 to give Eqn. 10, where ω_*N*_ is the ^15^N Larmor frequency (in radians per second):

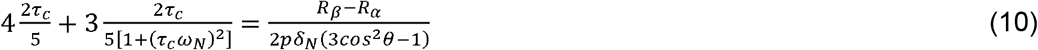

The right-hand side of Eqn. 10 is a constant once the relaxation rates have been measured. We therefore replace this side with the symbol ‘c’.

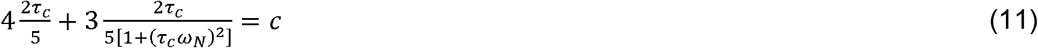

Solving Eqn. 11 for *τ*_*c*_ gives^20^,

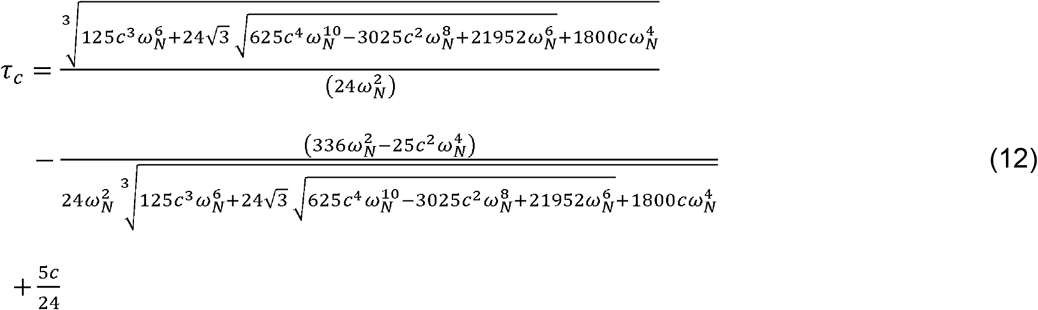

where,

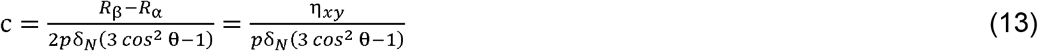

Using Eqns. 12 and 13, we recalculated *τ*_*c*_ = 19.8 and 22.5 ns, respectively, which are again approximately 6.5% less than the reported values. We concluded that Lee et al. contains an inadvertent error, and note that using a 700 MHz field (instead of the quoted 750 MHz) does reproduce the reported *τ*_*c*_ values.

We next surveyed the citing literature with the hypothesis that subsequent TRACT users may have similar miscalculations. As of early 2021, the original manuscript had been cited 120 times in PubMed. Half of all citing publications appeared in the years 2015-2020, indicating an increasing interest in this methodology 10-15 years after its’ original publication. We focused our analysis on manuscripts with sufficient data to validate calculations using our algebraic solution (Eq. 12). Table 1 shows that out of 120 manuscripts, three referenced the original TRACT paper without performing any cross-correlated relaxation experiments, six were methodological reviews, and seventy-eight used the TRACT methodology but did not provide enough experimental information to confirm their calculations. Thirty-three papers (30%) reported sufficient data to authenticate 65 total calculations.

**Table 1:** Published manuscripts that cite the TRACT paper^7^ categorized by: papers that reference the TRACT paper but provided no *τ*_*c*_ calculations; NMR methodological reviews that detail the TRACT experiment; papers that determined *τ*_*c*_ using the TRACT method but did not provide sufficient data for *τ*_*c*_ verification; and papers that did provide enough data for verification of *τ*_*c*_ calculations.

| **Paper Category** | **Number of Articles** | **Percentage of Articles (%)** |
| --- | --- | --- |
| Only Referenced | 3 | 2.5 |
| Reviews | 6 | 5.0 |
| Insufficient Data for Analysis | 78 | 65.0 |
| Sufficient Data for Analysis | 33 | 27.5 |

We defined an error measure by dividing the published *τ*_*c*_ by our algebraically determined value. For example, the original paper reported *τ*_*c*_ = 21 and 24 ns, while we calculated values of 19.8 and 22.5 ns, giving error ratios of 1.061 and 1.067, respectively. There are several features of this analysis worthy of report (Fig. 1A). First, 35% (23/65) of results are accurate (error ratio ≈ 1.0), clustering to within a 2% error interval of 0.99 and 1.01 and validating our algebraic approach. Second, 23% (15/65) of results cluster around 1.067 ± 0.03 (dotted line); this strongly suggests an error from the original TRACT report^11^ was propagated into the NMR literature. Finally, we note that 29% (19/65) of calculations have error ratios greater than 1.1 (>10% error). Extrapolating our results implies that ~65% of all citing literature (over 70 calculations) incorrectly estimate the rotational correlation time using the TRACT methodology. We also noted that researchers are more likely to overestimate the rotational correlation time, especially for low *τ*_*c*_ values. A scatter plot of *τ*_*c*_ versus error ratio demonstrates an inverse trend with highly erroneous values when *τ*_*c*_ < 2 ns (Fig. 1B). Plotting *τ*_*c*_ errors by year of publication underscores the persistence of these miscalculations in contemporary literature (Fig. 1C).

### Evaluation and analysis using experimental data

While the CCR experiments eliminate complications from chemical exchange and remote dipolar couplings (Eqns. 2 and 3), the existence of fast internal motions cannot be established from these data alone. Spin systems possessing non-negligible *τ*_*e*_ would artificially reduce *τ*_*c*_ values using Goldman’s relationship (Eqns. 6 and 12)^12^. This concern is especially relevant to the TRACT approach because spectra are often only collected in the directly-acquired ^1^H dimension, and then analyzed by integration over a chosen ^1^H_N_ region (typically ^1^H_N_ δ > 8 ppm) to improve S/N. This leads to significant signal overlap, especially in the high molecular weight target proteins for which these experiments were designed^10,11^. Overlap itself can be mitigated by acquiring 2D versions with indirect ^15^N evolution (although at a significant time expense), but confirmation of fast timescale motions still requires ^15^N{^1^H}-NOE data which are especially problematic in perdeuterated, high molecular weight systems^21^.

Our algebraic solution enables rapid point-by-point *τ*_*c*_ calculation, which we used to explore the signal overlap problem using [*U*-^15^N,^2^H]-OmpX prepared in DPC micelles (Fig. 2). The 1D ^1^H_N_ TROSY (i.e. N_α_ spin state) spectrum at a relaxation delay = 1 ms is well dispersed with high intensity from 8.5 – 8.0 ppm indicative of many overlapped spin systems and/or rapid local motions (Fig. 2A). As expected, regions with weak signal intensity (e.g. 9.8 ppm ≥ ^1^H δ ≤ 8.0 ppm) give wildly variable *τ*_*c*_ estimates (Fig. 2B). There is sufficient signal intensity between 9.5 – 8.8 ppm to calculate reasonable *τ*_*c*_ values of 30-50 ns. A significant drop in *τ*_*c*_ is observed as data approaches the center of the amide region (~8.5 ppm; Fig. 2C), which reflects OmpX’s unstructured loops 3 and 5^19^ that resonate at random coil chemical shifts^22^. Together, this demonstrates that arbitrary selection of a region for integration, without knowledge of underlying dynamic processes, is problematic. Even estimations from relatively invariable regions, such as 9.5 – 8.8 ppm, still possess high variance on a point-by-point basis. We applying a 200 point (5% of 4096 point amide region) sliding window as an optimal compromise for both sensitivity (via integration) and variability (region selection). The 200 points in the window are integrated and a *τ*_*c*_ prediction generated (Fig. 2C). As illustrated in Figure 2, the sliding window approach narrows the standard deviation for straight-forward identification of regions with consistent *τ*_*c*_ values. For example, calculations from 9.5 – 9.17 ppm, with and without a sliding window, result in *τ*_*c*_ of 37.92 ± 4.25 ns and 37.17 ± 0.59 ns, respectively (Fig. 2C,D). While commonly-applied assumptions about dispersed signals and global tumbling are useful when analyzing overlapped signals, point-by-point calculations coupled with the sliding window approach enable data-driven verification of consistent properties across the region of interest.

### Rigid-body approximation as source of systematic error

In its current form, the Goldman relation (Eqn. 6) and our algebraic solution (Eqn. 12) do not account for bond motions. As well-ordered protein regions generally possess backbone ^15^N_H_ 0.85 ≤ S^2^ ≤ 0.95^23,24^, we hypothesize the rigid-body assumption results in an underestimate of rotational correlation times estimated from TRACT experiments. We modified the spectral density function in Goldman’s relationship (Eqn. 6) to include an order parameter (Eqn. 14) and solved for τ as a function of B_0_, R_α_, R_β_, and S^2^ (Eqn. 15). Note, we use a simplified form of the model-free spectral density function^3^ that excludes fast timescale motions (*τ*_*e*_) which TRACT data alone is insufficient to estimate. Therefore, Eqns. 14 and 15 are only applicable to spin systems where *τ*_*e*_ has been established to be negligible through the process above.

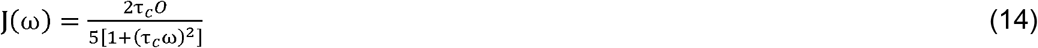

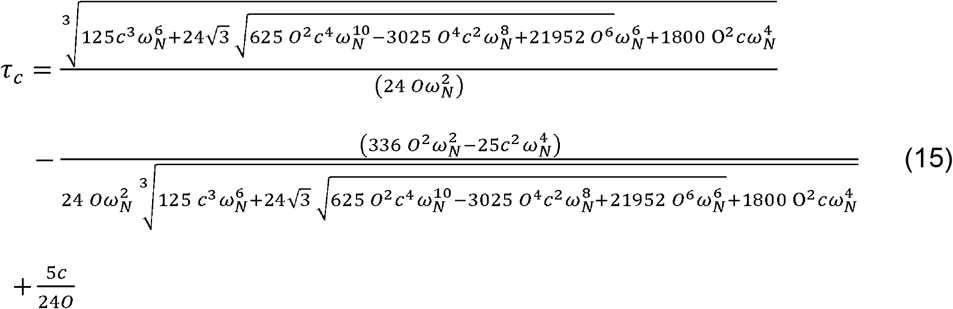

Where O is intended to stand for the S^2^ order parameter, and

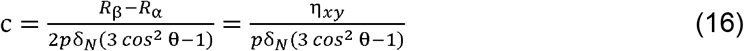

We then modelled the effect of S^2^ order parameters on *τ*_*c*_ estimation using Eqn. 15 (Fig. 3). Relative to the rigid body assumption, typical backbone order parameters (0.85 ≤ S^2^ ≤ 0.95) increase the rotational correlation time 5-25% depending on the CCR rate; this error increases to ≥45% at an S^2^ ≈ 0.7. Plotting errors in *τ*_*c*_ as a function of η_*xy*_ for various order parameters reveals a pronounced biphasic character centered ≈ 4 Hz that becomes largely invariant at η_*xy*_ > 10 ns (Fig. 3B). This simulation was performed with B_0_ = 800 MHz where η_*xy*_ = 4 Hz gives a *τ*_*c*_ of ~ 4 ns (assuming S^2^ = 1.0). There is insufficient data to estimate S^2^ from TRACT data alone; however, as shown, the rigid body assumption leads to significant underestimation of the rotational correlation time. This problem has been previously discussed by Wand and coworkers^25^ who found TRACT underestimated *τ*_*c*_ by an ≈ 20% for five different proteins, with errors ranging from 15-35%, when compared to more rigorous methods. Here we show that, in principle, these errors could be substantially-reduced by inclusion of an order parameter to the Goldman relationship.

### Potential sources of systematic error

The internuclear distance (*r*), difference of the two principal components of the axially-symmetric CSA tensor (*Δδ*_*N*_), and angle of the CSA tensor relative to the N-H bond vector (θ) are three additional parameters assumed constant in the equations above. These values are typically applied uniformly across the protein in ^15^N relaxation analyses although they’re well documented to be dependent on local structure^26,27^. Further, there are multiple commonly-employed values used throughout the literature, which are, themselves, interdependent and likely sources of systematic error^26,27^. For example, a static N-H *r* = 1.02 Å was used across NMR dynamics analyses until Ottiger and Bax calculated a vibrationally-corrected *r* = 1.041 ± 0.006 Å^28^. A much wider range of *Δδ*_*N*_ values have been reported, including −157 ± 19 ppm^29^, −172 ± 13 ppm^30^, and between −173.9 and −177.2 ppm^31^. Several studies demonstrate the CSA tensor is dependent on secondary structure^26,27,32^ with solid-state NMR experiments reporting average ^15^N CSA values = −187.9, −166.0, and −161.1 ppm for helices, strands, and turns, respectively; Ramamoorthy and colleagues go on to show that a change of 10^−2^ Å in N-H bond length or 1° deviation in θ could alter the calculated CSA tensor^27^. Recent values for θ include 15.7 ± 5° Fushman, 1998 #21}, 19.9°^31^, and 21.4 ± 2.3°^33^. To explore the effect of each parameter on the Goldman relationship, we calculated rotational correlation times as a function of each parameter individually and plotted the percentage error relative to *r* = 1.02 Å, θ = 17° and *Δδ*_*N*_ = −160 ppm (Fig. 4). While *r* and θ deviations generally effect rotational correlation time estimates within a ±5% error, variations of *Δδ*_*N*_ can result in errors up to ±15% (Fig. 4). It is important to note, however, that these errors approximate a mathematically-odd function centered around the elected value. That is, integration over many spin systems would cancel out small deviations around the average value for these three parameters, regardless of the parameter’s chosen magnitude. Although, this characteristic would have little benefit in situations when the geometric constant is obviously inappropriate, such as using a helical CSA tensor to evaluate a beta-stranded protein.

Finally, we consider the contribution of remote dipole-dipole interference with local CSA on measured transverse cross-correlated relaxation rates. These interactions are distinct from the *R*_*D*_ contributions in Eqns. 2 and 3 and do not cancel out when subtracting *R*_*β*_ from *R*_*α*_. Generally, remote protons are not found within 2.2 Å of a given ^1^H-^15^N spin system, regardless of secondary structure, due to steric exclusion^34^. Given the *r*^−3^, dependence of the dipole effect, protons at a distance of 2.2 Å contribute approximately 10 times less than protons at 1.02 Å to dipole relaxation. Liu and Prestegard have previously simulated the contributions of remote dipole/local CSA interference on measured CCR rates for the yARF1 protein^18^. They demonstrate that the average error in the CCR rate is 0.75% with an upper limit of ~3.5%{Liu, 2008 #17. In the perdeuterated case, which is essential for the study of proteins > 30 kDa, this effect would be even further minimized. Moreover, if the structure is known, the method described by Lui and Prestegard could be used to determine the error contribution.

## Conclusion

All NMR methods for estimating the rotational correlation time depend on a number of assumptions concerning how molecules behave in solution. The most comprehensive, and time-consuming, approach involves the collection of multidimensional T_2_, T_1_ and heteronuclear ^15^N{^1^H}-NOE experiments. Establishing which spin systems have suitable behavior is not trivial and difficult to automate; nonetheless, this strategy is independent of order parameters. This advantage is significant and enables site-specific *τ*_*c*_ estimation. Two major drawbacks are the difficulties associated with applying these experiments to high molecular weight systems, and the significant effect of remote dipolar couplings on measured longitudinal relaxation rates. In an attempt to circumvent complications from chemical exchange and remote protons, Lee and coworkers developed the TRACT experiment to estimate rotational correlation times from cross-correlated relaxation rates using the Goldman relationship.

Herein, we developed two algebraic solutions to the Goldman relationship for accurate calculations assuming the rigid-body approximation or a specific order parameter. These solutions enabled us to explore the boundaries of the Goldman relationship without relying on numerical minimization, which is computationally slow and potentially inaccurate. However, as we have discussed in this paper, accurate analysis of TRACT data also requires careful consideration. First, there is no way to directly detect spin systems with fast internal motions that would undervalue *τ*_*c*_ estimates. Second, experiments are frequently collected in a one-dimensional mode that is quite fast, but sensitive to signal overlap. The algebraic solutions facilitate rapid, point-by-point calculations for straightforward identification of appropriate spectral regions where global tumbling is likely to be dominant. Combining this approach with a sliding window simulates the advantages of integration while minimizing the inclusion of inappropriate spin systems. We also demonstrate that the rigid-body approximation can substantially underestimate TRACT-based rotational correlation time estimates. Our algebraic solution incorporates a simplified model-free spectral density function with order parameter that could, in principle, be set to an average backbone S^2^ ≈ 0.9 to further improve the accuracy of *τ*_*c*_ estimation. This has not been considered previously. Deviations in *r*, θ and *Δδ*_*N*_ contribute modest errors to *τ*_*c*_ estimation, although these would be expected to cancel out over a large number of spin systems. We hope our algebraic solutions and analytical strategies will increase the accuracy and application of the TRACT experiment.

## Acknowledgements

This study was supported by National Institutes of Health grant R00GM115814 (J.J.Z.), Indiana University start-ups (J.J.Z.), and the Indiana Precision Health Initiative (J.J.Z.). The 800 MHz NMR spectrometer used in this research was generously supported by a grant from the Lilly Endowment.

## Declarations

Funding: National Institutes of Health (R00GM115814 to J.J.Z.)

Conflicts of Interest/Competing interests: None

Availability of data and material: Available from Authors upon request

Code Availability: Published in SI and available at https://github.com/nomadiq/TRACT_analysis/blob/master/tract_algebraic_analysis.py

Authors’ contributions: SAR and JJZ developed the idea, analyzed the data, and wrote the paper. CD and HW prepared the OmpX sample and collected NMR data.

Ethics approval: N/A Consent to participate: N/A

Consent for publication: All of the authors have approved the contents of this paper and have agreed to the Journal of Biomolecular NMR’s submission policies.

