## Supplemental Information for "TRACT revisited: an algebraic solution for determining overall rotational correlation times from cross-correlated relaxation rates"

**Supporting Information**

**Listing S1:** Algebraic Solution for tau_c with and without order parameter S^2^

<https://github.com/nomadiq/TRACT_analysis/blob/master/tract.py>

```
import numpy as np


def default_tract_params():
    params = {"h": 6.62607004 * (1 / np.power(10, 34, dtype=np.longdouble))}
    params["mu_0"] = 1.25663706 * (
        1 / np.power(10, 6, dtype=np.longdouble)
    )  # vacuum permeability
    params["gamma_H"] = 267.52218744 * np.power(
        10, 6, dtype=np.longdouble
    )  # proton gyromagnetic ratio
    params["gamma_N"] = -27.116 * np.power(
        10, 6, dtype=np.longdouble
    )  # 15N gyromagnetic ratio
    params["r"] = 1.02 * (
        1 / np.power(10, 10, dtype=np.longdouble)
    )  # internuclear distance
    params["delta_dN"] = 160 * (
        1 / np.power(10, 6)
    )  # diff in axially symetric 15N CS tensor
    params["theta"] = 17 * np.pi / 180  # angle between CSA axis and N-H bond

    return params


def J(w, tc):
    return 0.4 * tc / (1 + (w ** 2 * tc ** 2))


def J_S2(w, tc, S2):
    return S2 * J(w, tc)


# function that returns evaluation of equation (12) with w_N and constant 'c' above as inputs
def tc_algebraic(Ra, Rb, field, params=None):

    if not params:  # use these defaults
        params = default_tract_params()

    h = params["h"]
    mu_0 = params["mu_0"]
    gamma_H = params["gamma_H"]
    gamma_N = params["gamma_N"]
    r = params["r"]
    delta_dN = params["delta_dN"]
    theta = params["theta"]

    B_0 = field * np.power(10, 6, dtype=np.longdouble) * 2 * np.pi / gamma_H  # in Tesla
    p = (
        mu_0
        * gamma_H
        * gamma_N
        * h
        / (16 * np.pi * np.pi * np.sqrt(2) * np.power(r, 3))
    )  # DD 1H-15N bond
    dN = gamma_N * B_0 * delta_dN / (3 * np.sqrt(2))  # 15N CSA
    w_N = B_0 * gamma_N  # 15N frequency (radians/s)

    c = (Rb - Ra) / (2 * dN * p * (3 * np.cos(theta) ** 2 - 1))

    t1 = (5 * c) / 24
    t2 = (336 * (w_N ** 2) - 25 * (c ** 2) * (w_N ** 4)) / (
        24
        * (w_N ** 2)
        * (
            1800 * c * (w_N ** 4)
            + 125 * (c ** 3) * (w_N ** 6)
            + 24
            * np.sqrt(3)
            * np.sqrt(
                21952 * (w_N ** 6)
                - 3025 * (c ** 2) * (w_N ** 8)
                + 625 * (c ** 4) * (w_N ** 10)
            )
        )
        ** (1 / 3)
    )
    t3 = (
        1800 * c * (w_N ** 4)
        + 125 * (c ** 3) * (w_N ** 6)
        + 24
        * np.sqrt(3)
        * np.sqrt(
            21952 * (w_N ** 6)
            - 3025 * (c ** 2) * (w_N ** 8)
            + 625 * (c ** 4) * (w_N ** 10)
        )
    ) ** (1 / 3) / (24 * w_N ** 2)

    return t1 - t2 + t3


# function that returns evaluation of equation (15) with w_N and constant 'c' above as inputs
def tc_algebraic_S2(Ra, Rb, field, S2, params=None):

    if not params:  # use these defaults
        params = default_tract_params()

    h = params["h"]
    mu_0 = params["mu_0"]
    gamma_H = params["gamma_H"]
    gamma_N = params["gamma_N"]
    r = params["r"]
    delta_dN = params["delta_dN"]
    theta = params["theta"]

    B_0 = field * np.power(10, 6, dtype=np.longdouble) * 2 * np.pi / gamma_H  # in Tesla
    p = (
        mu_0
        * gamma_H
        * gamma_N
        * h
        / (16 * np.pi * np.pi * np.sqrt(2) * np.power(r, 3))
    )  # DD 1H-15N bond
    dN = gamma_N * B_0 * delta_dN / (3 * np.sqrt(2))  # 15N CSA
    w_N = B_0 * gamma_N  # 15N frequency (radians/s)

    c = (Rb - Ra) / (2 * dN * p * (3 * np.cos(theta) ** 2 - 1))
    w = w_N

    t1 = (
        (
            125 * (c ** 3) * (w ** 6)
            + 24
            * np.sqrt(3)
            * np.sqrt(
                625 * (c ** 4) * (S2 ** 2) * (w ** 10)
                - 3025 * (c ** 2) * (S2 ** 4) * (w ** 8)
                + 21952 * (S2 ** 6) * (w ** 6)
            )
            + 1800 * c * (S2 ** 2) * (w ** 4)
        )
        ** (1 / 3)
    ) / (24 * S2 * w ** 2)

    t2 = (
        -1
        * (336 * (S2 ** 2) * (w ** 2) - 25 * (c ** 2) * (w ** 4))
        / (
            (
                24
                * S2
                * (w ** 2)
                * (
                    125 * (c ** 3) * (w ** 6)
                    + 24
                    * np.sqrt(3)
                    * np.sqrt(
                        625 * (c ** 4) * (S2 ** 2) * (w ** 10)
                        - 3025 * (c ** 2) * (S2 ** 4) * (w ** 8)
                        + 21952 * (S2 ** 6) * (w ** 6)
                    )
                    + 1800 * c * (S2 ** 2) * (w ** 4)
                )
                ** (1 / 3)
            )
        )
    )

    t3 = +(5 * c) / (24 * S2)

    return t1 + t2 + t3


# example

Rb = 64
Ra = 12
field = 750
S2 = 0.86

print(
    f"\n\u03C4_c = {tc_algebraic(Ra, Rb, field)*1000000000:.4f} ns where Rb = {Rb} Hz, Ra = {Ra} Hz, Field = {field} MHz"
)
print(
    f"\u03C4_c = {tc_algebraic_S2(Ra, Rb, field, S2)*1000000000:.4f} ns where Rb = {Rb} Hz, Ra = {Ra} Hz, Field = {field} MHz, S^2 = {S2}"
)

print("\n\nAdjusting \u03B8 to 20\n")
params = default_tract_params()
params["theta"] = 20 * np.pi / 180

print(
    f"\u03C4_c = {tc_algebraic(Ra, Rb, field, params)*1000000000:.4f} ns where Rb = {Rb} Hz, Ra = {Ra} Hz, Field = {field} MHz"
)
print(
    f"\u03C4_c = {tc_algebraic_S2(Ra, Rb, field, S2, params)*1000000000:.4f} ns where Rb = {Rb} Hz, Ra = {Ra} Hz, Field = {field} MHz, S^2 = {S2}"
)
```

**Listing S2:** Numerical Minimization Analysis

<https://github.com/nomadiq/TRACT_analysis/blob/master/tract_minimization_analysis.py>

```
import numpy as np
from scipy.optimize import minimize_scalar, minimize

# constants
h = 6.62607004 * (1/np.power(10,34, dtype=np.longdouble))  # Plank's
mu_0 = 1.25663706 * (1/np.power(10,6, dtype=np.longdouble)) # vacuum permeability
gamma_H = 267.52218744 * np.power(10,6, dtype=np.longdouble) # proton gyromagnetic ratio
gamma_N = -27.116 * np.power(10,6, dtype=np.longdouble)   # 15N gyromagnetic ratio
r = 1.02 * (1/np.power(10,10, dtype=np.longdouble))     # internuclear distance
delta_dN = 160 * (1/np.power(10, 6))             # axially symetric 15N CS tensor
theta = 17*np.pi/180                     # angle between CSA axis and NH bond

# field in MHz
field = 750

# derived field value in Tesla
B_0 = field * np.power(10, 6, dtype=np.longdouble) * 2 * np.pi / gamma_H # in Tesla

# equation 7
p = mu_0*gamma_H*gamma_N*h/(16*np.pi*np.pi*np.sqrt(2)*np.power(r,3))   # DD 1H-15N bond

# equation 8
dN = gamma_N*B_0*delta_dN/(3*np.sqrt(2)) # 15N CSA

w_N = B_0 * gamma_N           # 15N frequency (radians/s)

# measured relaxation rates
# Panel A Figure 4
Rb = 64 # Hz
Ra = 13 # Hz

#Panel B Figure 4
Rb = 80 # Hz
Ra = 22 # Hz

args = (w_N, Rb, Ra, p, dN, theta)

# Equation 15 function
def objective_function(tau_c, *args):

  #spectral density function
  def J(w_N, tau_c):
    return 0.4*tau_c/(1+(w_N**2*tau_c**2))
  #print(*args)
  (w_N, Rb, Ra, p, dN, theta) = args # unpack these constants inside the function
  return np.abs((4*J(0, tau_c) + 3*J(w_N, tau_c)) - ((Rb - Ra)/(2*p*dN*(3*np.cos(theta)**2-1))))

# guess tau_c
t = 10 * (1/np.power(10,8, dtype=np.longdouble)) # guess 10 ns
res = minimize_scalar(objective_function, args=args, method='Brent')
print(f'Given: field = {field} MHz, Rb = {Rb} Hz and Ra = {Ra} Hz')
print(f'tau_c: {res.x} seconds -- Using Brent Minimization')
print()
res = minimize(objective_function, t, args=args, method='BFGS')
print(f'Given: field = {field} MHz, Rb = {Rb} Hz and Ra = {Ra} Hz')
print(f'tau_c: {res.x} seconds -- Using BFGS Minimization')
print()
res = minimize(objective_function, t, args=args, method='Powell')
print(f'Given: field = {field} MHz, Rb = {Rb} Hz and Ra = {Ra} Hz')
print(f'tau_c: {res.x} seconds -- Using Powell Minimization')
print()
res = minimize(objective_function, t, args=args, method='TNC')
print(f'Given: field = {field} MHz, Rb = {Rb} Hz and Ra = {Ra} Hz')
print(f'tau_c: {res.x} seconds -- Using TNC Minimization')
print()
```
